# Characterization of a crustacean hyperglycemic hormone of the horsehair crab *Erimacrus isenbeckii*

**DOI:** 10.1101/2025.04.26.650751

**Authors:** Kenji Toyota, Asami Kajimoto, Yushi Ando, Ken Takeuchi, Tsuyoshi Ohira

## Abstract

The horsehair crab *Erimacrus isenbeckii* is widely distributed from Alaska and the Bering Sea through Southern Sakhalin, and in Japan from the coastal areas of Hokkaido to the Sea of Japan. In the Hokkaido area, although strict resource management has been promoted by setting an allowable catch limit, the catch amount has remained at a low level. While knowledge on larval rearing methods is accumulating in relation to seedling production techniques, information on adult growth and molting is limited, due to a deep-sea species that requires a long period for growth. In decapod crustaceans, the sinus-gland/X-organ complex in the eyestalk ganglion synthesizes and secrets various neuropeptides such as crustacean hyperglycemic hormone (CHH) to regulate the homeostasis of blood glucose levels. In this study, combined isolation of sinus gland peptides by reverse-phase high-performance liquid chromatography (RP-HPLC) and amino acid sequencing, and transcriptome analyses using male and female eyestalk ganglion has been successfully identified EiCHHa. The *in vivo* assays of EiCHHa using the blue swimming crab *Portunus pelagicus* revealed EiCHHa has a hyperglycemic effect, as well as CHHs in other decapod crustaceans. Additionally, we successfully demonstrated the sexual differences in the transcriptomic profiles between males and females. Especially, two sinus gland-derived neuropeptides (EiCHHb and a crustacean female sex hormone (EiCFSH)) were isolated as female-biased transcripts, suggesting that both hormones may have female-specific roles such as the development of female characteristics and reproduction.

## 1. Introduction

The horsehair crab *Erimacrus isenbeckii* is widely distributed from Alaska and the Bering Sea through Southern Sakhalin, and in Japan from the coastal areas of Hokkaido to the Sea of Japan (Jinbo et al., 2005). In Japan, it is mainly distributed in the coastal areas of Hokkaido, especially in the Sea of Okhotsk and the Eastern Pacific Ocean (e.g., Funka Bay), and is an important target species for fisheries due to its delicious taste and high price. In Hokkaido, strict resource management has been promoted by setting an allowable catch limit, but the catch has remained at a low level, and active stock enhancement through the release of seedlings is desired. Several organizations have developed seedling production technology for this species, but the survival rate of larvae is unstable (Ichikawa et al., 2018; Jinbo et al., 2005), and therefore stable seedling production technology has not yet been established. While knowledge on larval rearing is accumulating in relation to seedling production techniques, information on adult growth and molting are limited, due to a deep-sea species that requires a long period for growth (Abe 1982). Based on these situations, no physiological research has been done using the horsehair crab.

In decapod crustaceans, the sinus-gland/X-organ complex in the eyestalk ganglion synthesizes and secrets various neuropeptides such as crustacean hyperglycemic hormone (CHH)-superfamily including the CHH, molt-inhibiting hormone (MIH), vitellogenesis-inhibiting hormone (VIH) to regulate the homeostasis of blood glucose levels, molting, and ovarian maturation (Katayama et al., 2013). Generally, the amounts and profiles of each neuropeptide varied by species, molting period, and breeding period (Green et a., 2023; Zmora and Chung, 2014). Indeed, crab species are able to predict a neuropeptide profile as two structural isoforms of CHH and a single MIH, referred to in the green crab *Carcinus maenas, Callinectes sapidus*, and red deep-sea crab *Chaceon quinquedens* (Chung and Webster, 1996; Green et a., 2023; Zmora and Chung, 2014). Moreover, transcriptome has enabled the construction of comprehensive gene sets. In terms of eyestalk-derived neuropeptide discovery, this tool has been applied and successfully identified various putative neuropeptides in several crab species such as Chinese mitten crab *Eriocheir sinensis* (Xu et al. 2015; Pang et al. 2019), hydrothermal vent crab *Austinograea alayseae* (Hui et al., 2017), the swimming crab *Portunus trituberculatus* (Lv et al., 2017). Our recent transcriptome revealed that transcriptomic profiles were different between females and males in the kuruma prawn *Marsupenaeus japonicus* (Toyota et al., 2023a). In this study, we conducted the eyestalk ganglion RNA-seq analysis and revealed the repertoire of genes encoding the eyestalk-derived neuropeptides including some CHH-family molecules in adult males and females *E. isenbeckii*. Then, one of the CHH-family peptides was purified by RP-HPLC from the sinus glands, and finally, its hyperglycemic activity was validated.

## 2. Materials and methods

### 2.1. Ethical statement

Fishing permission for horsehair crabs for this study was granted by the Hokkaido Governor (Hokkaido, Japan). All efforts were made to minimize the suffering of the animals.

### 2.2. Purification of CHH-family peptides from *E. isenbeckii*

Peptides were extracted from the 10 sinus glands of mature males and females using the same methods described previously (**Figure 1**; Yang et al. 1995). The extract was subjected to a Sep-Pak C18 Cartridge (Waters, Milford, MA, USA), which was eluted with 60 % acetonitrile in 0.05 % trifluoroacetic acid (TFA). After concentration, the resultant solution was applied to reversed-phase high-performance liquid chromatography (RP-HPLC) on a Shodex Asahipak ODP-50-4E column (4.6 × 250 mm, Showa Denko, Tokyo, Japan). Elution was performed with a 40-min linear gradient of 10–60 % acetonitrile in 0.05 % TFA, 1-min linear gradient of 60-80% acetonitrile in 0.05% TFA, and 5-min holding at 80% acetonitrile in 0.05% TFA at a 1 ml/min flow rate. Elution was monitored at 225 nm, and each peak fraction was collected manually.

**Figure 1.**
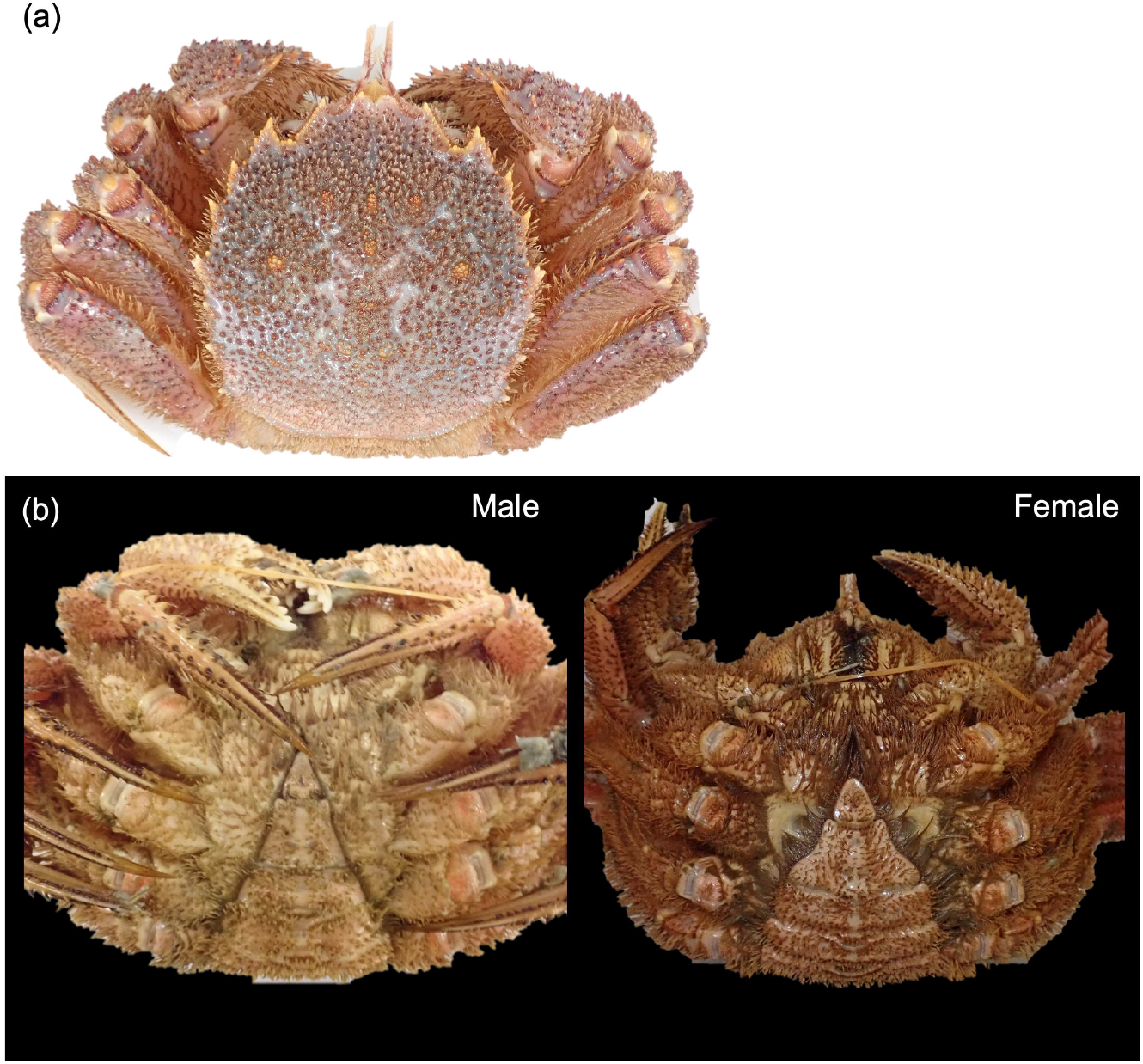
Photo images of horsehair crab. Dorsal view (a). Ventral views of male (left) and female (right) (b). The difference in abdominal shapes was used to distinguish males and females.

### 2.3. Mass spectral and amino acid sequence analyses

Mass spectra of the purified CHH (EiCHHa) was measured on a MALDI-TOF mass spectrometer (AXIMA®-CFR, Shimadzu, Kyoto, Japan) with α-cyano-4-hydroxycinnamic acid as a matrix in the positive ion mode. N-terminal amino acid sequences of the two CHH-family peptides were analyzed on PPSQ 51A protein sequencers (Shimadzu).

### 2.4. Pyroglutamate aminopeptidase digestion

The purified EiCHHa was dissolved in 50 mM sodium phosphate buffer (pH 7.0) containing 1 mM EDTA and 10 mM dithiothreitol (DTT), and then its N-terminal pyroglutamate was digested with 0.2 mU of *Pfu* pyroglutamate aminopeptidase (Takara Bio, Otsu, Japan) at 50°C for 12 h. After the initial incubation, the same amount of enzyme was added and further reacted for 12 hours. After the second reaction, a half volume of 1 M HCl was added to stop enzymatic digestion. The reaction product was applied to RP-HPLC under the same conditions described above.

### 2.5. Sample preparation for RNAseq

In April 2020, female and male horsehair crabs were collected from Funka Bay in Hokkaido, Japan **(Figure 1a)**. Four healthy males (wet weight: 409.75 ± 48.85 g) and females with no brooding (wet weight: 293.75 ± 29.44 g) were used for RNAseq analysis **(Figure 1b)**.

Both sides of the eyestalks were cut, and then each eyestalk ganglion, including the sinus gland, was extracted **(Figure 1c)**. Each sample (a total of 16 samples consisting of four left- or right-eyestalks) was stored separately in RNAlater reagent (Thermo Fisher Scientific, Waltham, MA, USA) until use. Total RNA was extracted using ISOGEN II (Nippon Gene Co Ltd., Tokyo, Japan) and the RNeasy Micro Kit (Qiagen, Valencia, CA, USA) from the right-side eyestalk ganglion, following the manufacturer’s protocols. The quality and concentration of total RNA were validated by NanoDrop (Thermo Fisher Scientific) and 2100 Bioanalyzer (Agilent Technologies, Santa Clara, CA, USA). All right-side eyestalk ganglion samples were used for the *de novo* transcriptome.

### 2.6. Construction of cDNA libraries, sequencing, *de novo* assembly, and differential expression analysis

The cDNA libraries were constructed using TruSeq stranded mRNA Kit (Illumina, San Diego, CA, USA) by Macrogen service (Macrogen Japan, Kyoto, Japan). All libraries were sequenced using an Illumina HiSeq2500 platform that was equipped with a 101 bp paired-end module. The quality of output sequences was inspected using the FastQC program (version 0.11.2). All reads from the female or male group (each of four biological replicates) were assembled together using the RNAseq *de novo* assembler Trinity (version 2.9.1) with the paired-end mode (Grabherr et al., 2011). Then female- and male-assembled transcriptomes were merged as a single fasta file and then submitted to the EvidentialGene tr2aacds pipeline (available online at: https://sourceforge.net/projects/evidentialgene/) for generating a single assembly with minimal redundancy while maximizing the maintenance of long coding sequence regions in each contig. Finally, the tr2aacds pipeline produced the “okay” set of transcripts that were regarded as optimal and representative transcript datasets that were used as a reference transcriptome for the mapping process. The reads from each biological replicate were mapped to the assembled transcripts for quantification by Salmon (version 1.1.0) with the “--dumpEq” option (Patro et al., 2017). Then, contigs were clustered based on the proportion of shared reads and expressions by Corset (version 1.09) (Davidson and Oshlack, 2014). Corset generated the clusters and outputs as a table of counts containing the number of reads uniquely assigned to each cluster. The completeness of orthologs of the Corset-generated transcriptome was examined using BUSCO version 5.4.3 against eukaryote_odb10 (Creation date: 2020-09-10, number of species: 70, number of BUSCOs: 255).

Corset generated the clusters and outputs a table of counts containing the number of reads uniquely assigned to each cluster. Using BLASTX (threshold E-values = 1E-03) by AC-DIAMOND package (version 0.9.22) (Mai et al., 2018), Corset-assembled cluster’s transcripts were aligned with the NCBI protein database NR (non-redundant). By using the Corset-generated count data, differentially expressed genes between females and males were calculated using the DESeq2 package in the SARTools package (version 1.6.6) between females and males (Varet et al., 2016). Principal component analysis, hierarchical clustering, and MA-plot were also conducted by the DESeq2 package in the SARTools package.

For gene ontology (GO) and Kyoto Encyclopedia of Genes and Genomes (KEGG) enrichment analyses using DEGs, Corset-assembled cluster’s transcripts were re-aligned with the protein database of the swimming crab *Portunus trituberculatus*, (Portunus_trituberculatus_gca017591435v1.GCA017591435v1.pep.all.fa) (Tang et al., 2020). Then, GO and KEGG enrichment analyses were conducted by DAVID browser (version 2021, last accessed: 25th February 2023).

### 2.7. Multiple sequence alignments and phylogenetic analysis

Protein sequences of CFSHs and CHH superfamily peptides were retrieved from online databases **(Table S1)**. Multiple alignments of the amino acid sequences were constructed using ClustalW (https://www.genome.jp/tools-bin/clustalw; last access by 2023 April 7). Phylogenetic analysis was conducted on their amino acid sequences using the Neighbor-joining framework of MEGA 11 software (Tamura et al., 2021). Bootstrap analysis, including 1,000 bootstrap replications, was applied to assess confidence in nodal support.

### 2.8. Cloning of *EiCHHa* gene from cDNA

Total RNAs were extracted from eyestalks using ISOGEN II. First-strand cDNA was synthesized using 250 ng of total RNA using SuperScript III with oligo dT primer (Life Technologies, Carlsbad, CA, USA) according to the manufacturer’s protocol, and then store at -20□ until use. A decoded amino acid sequence of putative EiCHHa was used as a query, and a homology search was performed against a gene catalog with a d*e novo* assembly. Then, only one transcript was retrieved with high similarity including 5’ and 3’ untranslated regions (Cluster-6453.16696). Based on this, the primers were designed for cDNA cloning as follows: forward and reverse primer sequences are “5’-GAGAGAGCCAGTATATA-3’ “and “5’-GTTCATCTTCCGGAGTGGAG-3’ “, respectively. The PCR was conducted using TaKaRa Ex Taq^®^ Hot Start Version (Takara Bio), and its PCR parameters were as follows: initial denaturation (94 °C for 3 min), followed by 45 cycles of 94°C for 30s, 50°C for 30s, 72°C for 2 min, and a final extension at 72 °C for 3 min. PCR amplicons were subcloned using a Mighty TA-cloning kit (Takara Bio) and those nucleotide sequences were analyzed.

### 2.9. Bioassay for hyperglycemic activity

Sixteen male individuals of horsehair crab were purchased from a fish market and acclimated for 24 hours in a 4°C recirculating seawater tank at the Tokyo University of Science (Oshamambe Campus: Oshamambe Town, Hokkaido, Japan). Subsequently, 8 individuals underwent bilateral eyestalk ablation, while the remaining 8 individuals were used as the control group (without eyestalk ablation) and maintained under fasting conditions. Blood samples (100 µL) were collected daily until day 6, and their blood glucose levels were measured with the glucose-oxidase peroxidase method (Yang et al. 1995).

Instead of *E. isenbeckii*,bioassays were conducted by using the red swamp crayfish *Procambarus clarkii* and the blue swimming crab *Portunus pelagicus*. Adult *P. clarkii* (carapace width: about 5.0 cm) and *P. pelagicus* (cephalothorax length: 3.5–4.0 cm), consisting of both sexes, were used in this study. These *P. clarkii* were collected from an irrigation canal in Hiratsuka City, Kanagawa Prefecture, Japan. These *P. pelagicus* were purchased from a fish store (Mie Prefecture, Japan). Both species were maintained at room temperature throughout the experiment. Bilateral eyestalk ablation was performed, followed by fasting at room temperature for two days prior to injection. Purified EiCHHa (1.0 μg/individual) in 100 μl of the physiological saline (SA) was injected into *P. clarkii* or *P. pelagicus* that had been eyestalk ablated 2 days earlier (n = 6 for *P. clarkii*; n = 5 for *P. pelagicus*). Hemolymph (100 μl each) was taken just before injection and 2 h after injection. SA was used as a negative control (n = 6 for *P. clarkii*; n = 5 for *P. pelagicus*). Hemolymph glucose levels were determined with the glucose-oxidase peroxidase method (Yang et al. 1995). The hyperglycemic activity was analyzed by Welch’s t-test using R (version 4.2.2) (R Core Team 2022).

## 3. Results

### 3.1. Purification and N-terminal amino acid sequences from *E. isenbeckii* CHH peptide

The sinus gland-derived peptide’s profiles of males and females were obtained from a total of 10 sinus glands by RP-HPLC **(Figure 2)**. In the TOF-MS spectra of the highest peak fractions in RP-HPLC chromatograms **(Figure 2A and 2B)**, protonated molecular ion peaks of males and females were observed at m/z 8387 and 8390, respectively. Therefore, these peak fractions were combined, and the contained peptide was subjected to N-terminal amino acid sequence analysis, but no sequence data was obtained. These results, along with previous reports on the amino acid sequences of crab CHHs (Katayama et al. 2013), strongly suggested that the N-termini of this peptide was blocked by N-terminal pyroglutamylation. Therefore, this peptide was digested with *Pfu* pyroglutamate aminopeptidase. The protein sequencing of the deblocked peptide demonstrated that partial N-terminal amino acid sequence was “IYDTSCKGVYDRGLFSDLEHVCDDCYNLYRNSHVAS”, and it is decided as CHH molecule of *E. isenbeckii* (EiCHHa).

**Figure 2.**
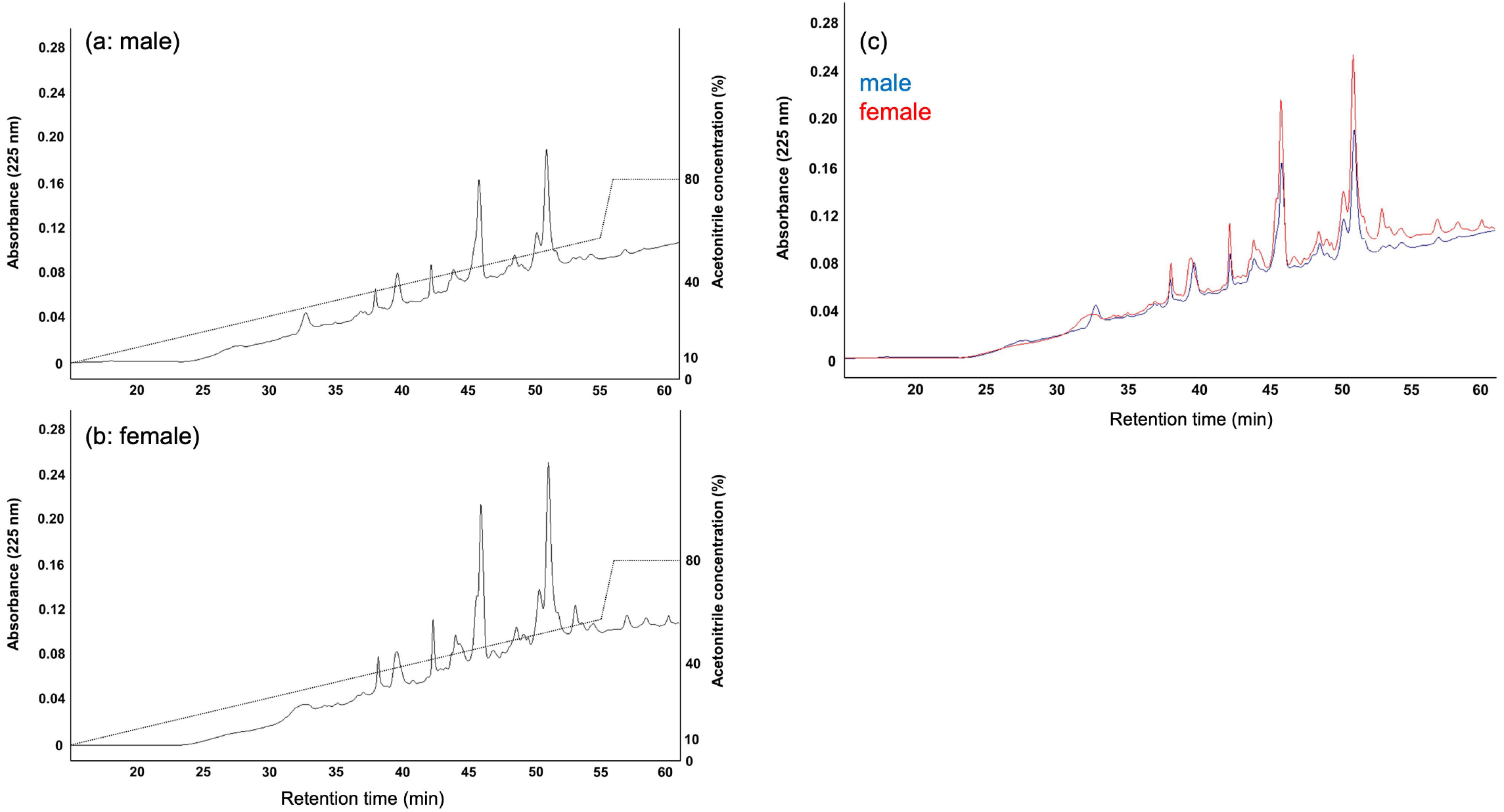
RP-HPLC elution profile of extracts of 10 sinus glands of females (a), males (b), and both sexes (c) of *E. isenbeckii*. Chromatographic conditions were described in “Materials and methods”. Arrows indicate the target fractions. The concentration of acetonitrile was indicated by the dotted line.

### 3.2. Sequence assembly, annotation, and extraction of CHH family genes

The *de novo* transcriptome assembly processes produced 54,183 putative transcripts using an analytical pipeline with Trinity, Evidentialgene, and Corset. Among them, 17,272 transcripts had significant BLAST similarity hits with publicly available protein sequences. The final transcripts were evaluated by BUSCO and obtained 96.9% of for completeness based on expectations of gene content against the eukaryotic gene dataset, indicating that our *de novo* assembly pipeline built reliable comprehensive gene models **(Table 1)**. Principal component analysis and clustering of the entire gene expression dataset indicated that the transcriptome profiles were well separated between female and male samples **(Figure S1a-c)**.

**Table 1.** Transcript completeness from BUSCO analysis.

| BUSCO evaluation metrics | Result |
| --- | --- |
| Complete BUSCOs (C) | 247 (96.9%) |
| Complete and single-copy BUSCOs (S) | 218 (85.5) |
| Complete and duplicated BUSCOs (D) | 29 (11.4) |
| Fragmented BUSCOs (F) | 6 (2.4%) |
| Missing BUSCOs (M) | 2 (0.7%) |
| Total BUSCO groups searched | 255 |

Our transcriptome successfully estimated the general sinus gland-derived neuropeptides as follows: three transcripts of crustacean hyperglycemic hormones (EiCHHa, EiCHHb, EiCHHc), molt-inhibiting hormone (EiMIH), red pigment-concentrating hormone (RPCH), two transcripts of pigment dispersing hormone (PDH), and crustacean female sex hormone (EiCFSH) **(Table 2)**. In terms of CHH, three transcripts of CHH (Cluster-6453.16696: EiCHHa; Cluster-6453.19920: EiCHHb, and Cluster-6453.24714: EiCHHc) were predicted **(Table 2)**.

**Table 2.**
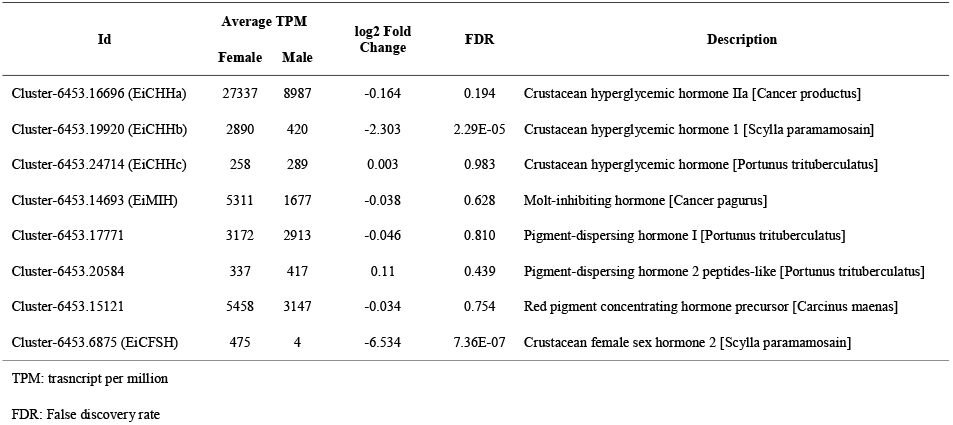
List of transcripts encoding sinus gland-derived peptides.

### Retrieve and cloning of the *EiCHHa*

Based on this amino acid sequence used as a query, high-homologous transcripts were surveyed in the assembled transcriptome, resulting in a single transcript Cluster-6453.16696 being successfully retrieved. When constructing a phylogenetic tree using EiMIH as an outgroup and three EiCHHs, all EiCHHs formed a group with CHHs from decapods **(Figure 3)**. EiCHHa was expressed highest in both male and female eyestalks among all CHH transcripts **(Table 2)**. Based on the assembled sequence, its full-length sequence consisted of 615 bp including an open reading frame (ORF) (567 bp), and 3’-UTR (48 bp). The ORF was conceptually translated into a putative prepropeptide comprising 189 amino acid residues **(Figure 4)**. The translated amino acid sequences showed a 100% identity with the peptide sequence Cluster-6453.16696 **(Figure 4)**. This gene was named as EiCHHa. The deduced amino acid sequences of EiCHHa showed the typical CHH structures, consisting of a signal peptide, CHH precursor-related peptide, and conserved residues (e.g., KR as dibasic cleavage site and GKKK as amidation signal sequence) **(Figure 4)**.

**Figure 3.**
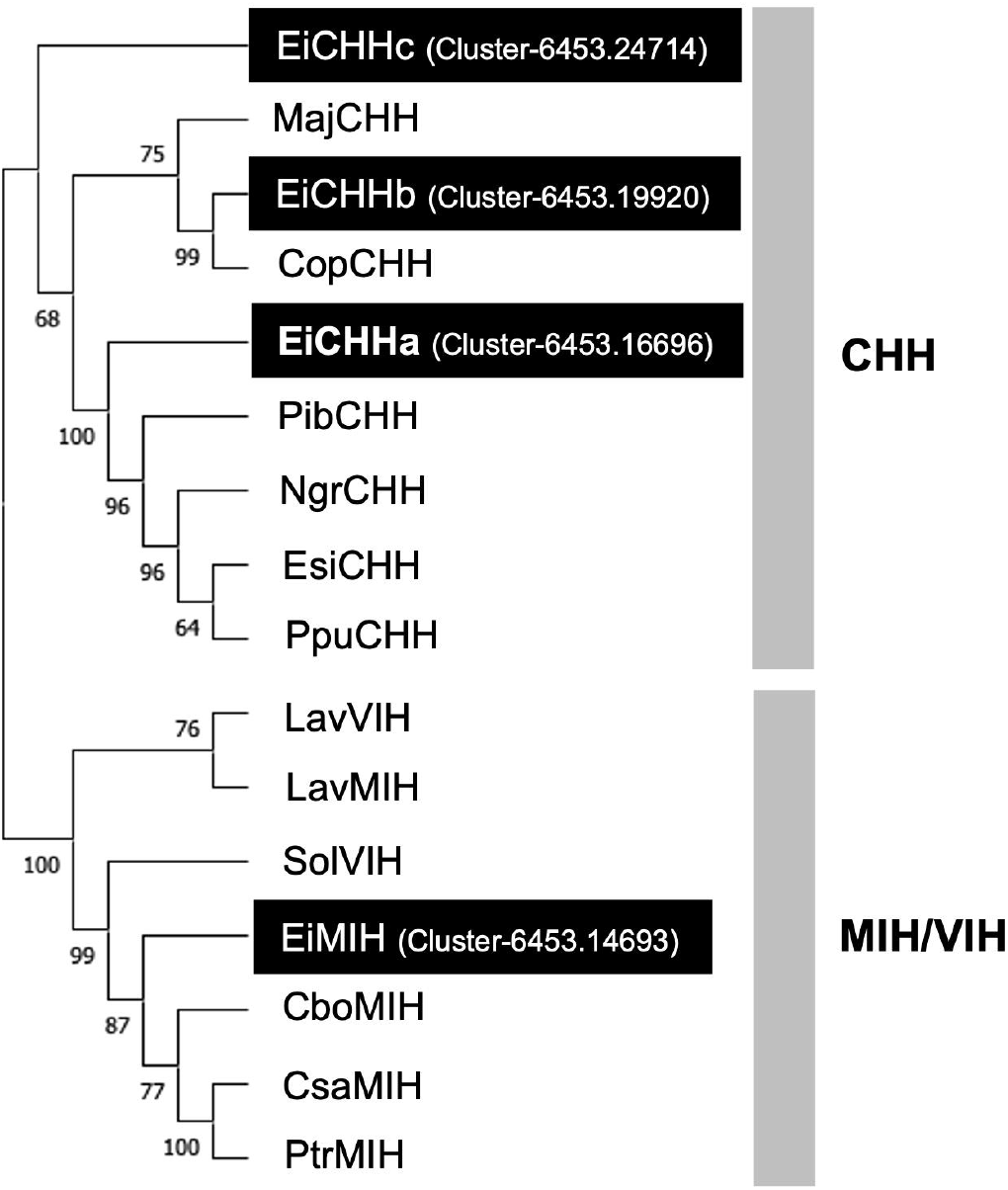
Maximum likelihood tree of the amino acid sequences of CHH superfamily with 1,000 bootstraps among crustaceans. Accession numbers are in **Table S1**.

**Figure 4.**
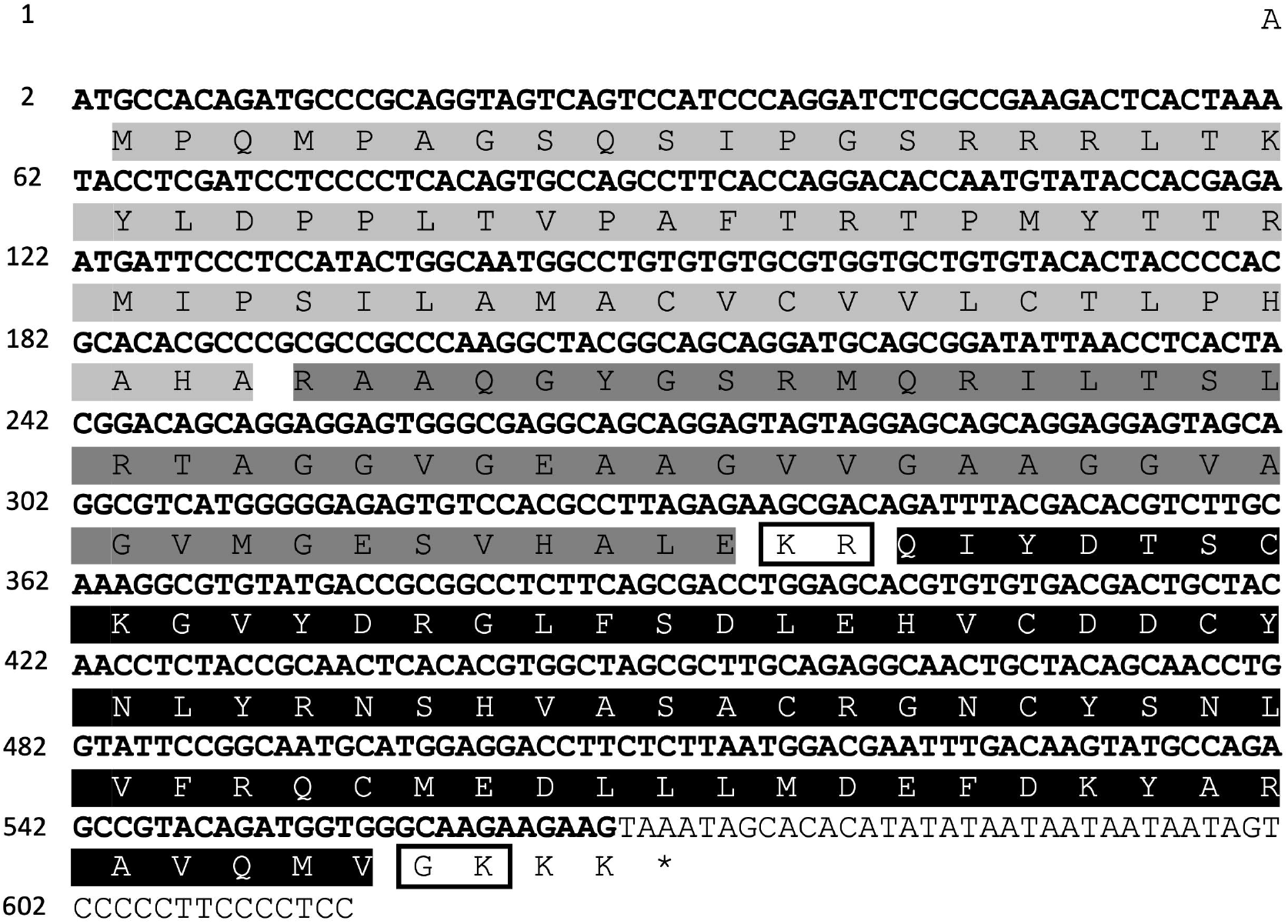
Nucleotide and deduced amino acid sequences of a cDNA encoding the EiCHHa precursor. The amino acid sequence of the putative signal peptide is marked by light gray, the putative CHH precursor-related peptide is marked by dark gray, and the putative mature EiCHHa is marked by black. The KR residues with an open box indicate the putative dibasic cleavage site. The GK residues with an open box indicate the putative amidation signal.

**Figure 5.**
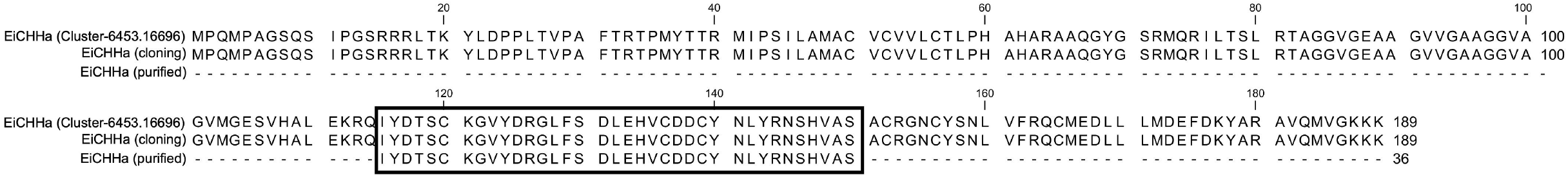
Alignment of EiCHHa with deduced amino acid sequences by de novo assembly, cDNA cloning, and purification from sinus gland extraction.

### Validation of hyperglycemic activity of EiCHHa using the red swamp crayfish and the blue swimming crab

To investigate whether blood glucose levels decrease in adult *E. isenbeckii* male under bilaterally eyestalk ablation conditions, blood glucose levels were monitored. The results showed that, although blood glucose levels decreased for 6 days fasting, there was no rapid decrease caused by eyestalk ablation, and no significant differences were observed compared to the control group without eyestalk ablation (Figure S3). From these results, it was concluded that evaluating the hyperglycemic effects of EiCHHa using *E. isenbeckii* is challenging. Instead, the red swamp crayfish *P. clarkii* and the blue swimming crab *P. pelagicus* were used for *in vivo* experiments.

Glucose levels in the hemolymph of *P. pelagicus* that were injected with the saline at 2 days after bilateral eyestalk ablation were 13.0 ± 8.4 μg/ml (mean ± SD, n = 5). On the other hand, EiCHHa-injected *P. pelagicus* with bilateral eyestalk ablation showed increased hemolymph glucose levels at 612.1 ± 104.9 μg/ml (n = 5) **(Figure 6a)**. On the other hand, EiCHHa-injected *P. clarkii* with bilateral eyestalk ablation showed no significant increased hemolymph glucose levels at 9.8 ± 57.0 μg/ml (n = 6) compared to saline injected group (- 18.1 ± 114.3 μg/ml, n = 6) **(Figure 6b)**.

**Figure 6.**
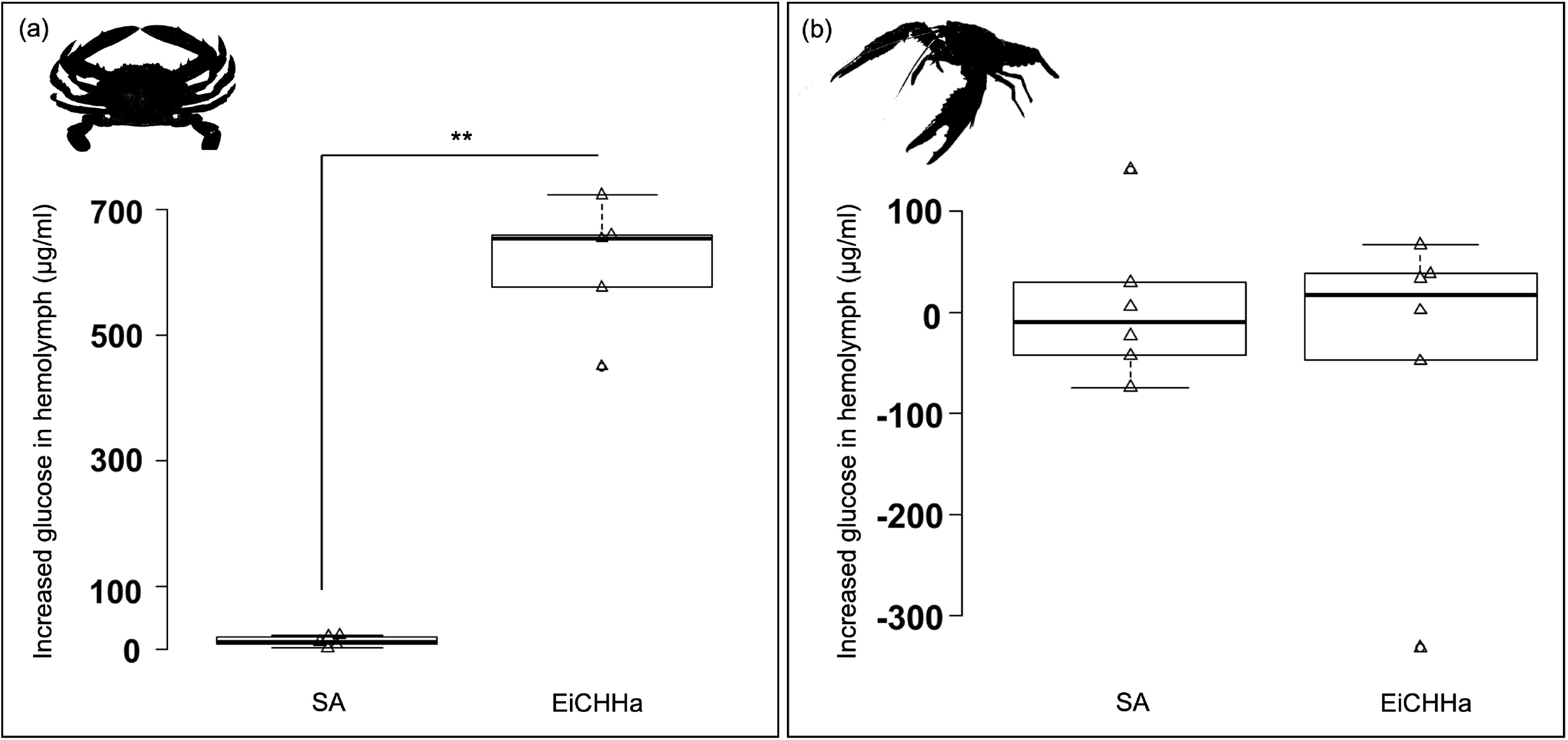
The hyperglycemic activity of EiCHHa using swimming blue crab *Portunus pelagicus* (a) and red swamp crayfish *Procambarus clarkii* (B). SA indicates the saline used as a negative control. The *p*-value was calculated by Welch’s t-test (**: *p* < 0.01).

### Profiles of male- and female-biased transcriptomes

Totally, 203 male- and 649 female-biased differential expressed transcripts (DETs) were screened with a false discovery rate < 0.05 **(Figure S1d)**. However, 105 (51%) and 202 (31%) DETs were categorized into functionally unknown groups in males and females, respectively. Of the top ten male-biased transcripts, six were cuticle-related transcripts, while no cuticle-related DET was found in the female-biased DET list **(Table 3)**. In terms of female-biased DEGs, EiCFSH and EiCHHb (Cluster-6453.19920) were selected as the top 10 DEGs **(Table 3)**. The phylogenetic tree of CFSH suggests an initial gene duplication giving rise to two types (CFSH1 and CFSH2), EiCFSH was categorized into CFSH1 group **(Figure S3)**. Moreover, our transcriptome revealed that EiCHHb (Cluster-6453.19920) expression level was significantly higher in females than in males, but not in other EiCHHc (Cluster- 6453.24714) **(Table 1)**. Although a large dispersion of expression profiles among individuals, the expression level of EiCHHa seemed to be higher in females as well **(Table 1)**. Besides, KEGG enrichment analysis provided four pathways as follows: “Oxidative phosphorylation”, “Ribosome”, “Citrate cycle (TCA cycle)”, and “Metabolic pathways” **(Table 4)**, suggesting that energy metabolism-related pathways may activate in female eyestalk ganglion.

**Table 3.** List of top 10 male- and female-biased expressed transcripts.

| Id | Average TPM |  | log2<br>Fold<br>Change | FDR | Description |
| --- | --- | --- | --- | --- | --- |
|  | Female | Male |  |  |  |
| Male-biased |  |  |  |  |  |
| Cluster-6453.39993 | 0 | 547 | 9.9 | 2.355E-05 | Cuticle protein [Cancer pagurus] |
| Cluster-6453.39918 | 2 | 3019 | 9.7 | 1.383E-05 | Cuticle proprotein proCP2.1 [Callinectes sapidus] |
| Cluster-6453.39959 | 10 | 3636 | 8.0 | 3.232E-06 | Cuticle protein CUT8 [Portunus pelagicus] |
| Cluster-6453.40043 | 0 | 273 | 7.6 | 4.068E-04 | Cuticle protein CUT12 [Portunus pelagicus] |
| Cluster-62.0 | 0 | 40 | 5.3 | 8.331E-04 | Cuticle protein AMP1A-like [Penaeus vannamei] |
| Cluster-6453.40449 | 5 | 185 | 4.8 | 3.820E-05 | Arthrodial cuticle protein AMP13.4 [Callinectes sapidus] |
| Cluster-6453.1332 | 7 | 182 | 4.5 | 8.122E-08 | Cuticle protein AM/CP1114 [Portunus trituberculatus] |
| Cluster-6453.39934 | 2 | 543 | 2.6 | 3.781E-03 | Cytochrome c1-like [Hyalella azteca] |
| Cluster-6453.26523 | 71 | 277 | 1.8 | 1.643E-07 | Putative arrestin-like [Penaeus vannamei] |
| Cluster-6453.12224 | 64 | 798 | 1.6 | 3.914E-03 | Crustin 2, partial [Portunus trituberculatus] |
| **Female-biased** |  |  |  |  |  |
| Cluster-6453.6875 (EiCFSH) | 475 | 4 | -6.5 | 7.356E-07 | Crustacean female sex hormone 2 [Scylla paramamosain] |
| Cluster-6453.34896 | 36 | 1 | -5.7 | 4.311E-05 | Sulfotransferase 1C4-like isoform X5 [Penaeus vannamei] |
| Cluster-6453.6829 | 173 | 10 | -4.1 | 5.943E-22 | Putative serine/threonine-protein kinase 2 [Portunus trituberculatus] |
| Cluster-6453.15450 | 1676 | 96 | -4.0 | 2.563E-15 | Triosephosphate isomerase B [Portunus trituberculatus] |
| Cluster-6453.18007 | 7514 | 1358 | -2.4 | 1.063E-23 | Serine protease 4 [Scylla paramamosain] |
| Cluster-6453.19920 (EiCHHb) | 2890 | 420 | -2.3 | 2.291E-05 | Crustacean hyperglycemic hormone 1 [Scylla paramamosain] |
| Cluster-6453.17557 | 4362 | 956 | -2.0 | 2.622E-12 | Leucine-rich repeat neuronal protein 3-like [Penaeus vannamei] |
| Cluster-6453.20206 | 137 | 30 | -2.0 | 2.658E-07 | Serine protease 4 [Scylla paramamosain] |
| Cluster-6453.17457 | 17130 | 4109 | -1.9 | 9.683E-12 | Transmembrane protein 205-like [Penaeus vannamei] |
| Cluster-6453.25344 | 680 | 171 | -1.8 | 5.550E-07 | Monocarboxylate transporter 14-like [Penaeus vannamei] |

TPM: trasncript per million
FDR: False discovery rate

**Table 4.** List of enriched KEGG pathways in female-biased expressed genes.

| KEGG pathway terms | Count | % | Pop Hits | Pop Total | PValue | FDR |
| --- | --- | --- | --- | --- | --- | --- |
| Oxidative phosphorylation | 18 | 4.5 | 103 | 3047 | 4.63E-07 | 3.38E-05 |
| Ribosome | 17 | 4.2 | 113 | 3047 | 8.44E-06 | 3.08E-04 |
| Citrate cycle (TCA cycle) | 6 | 1.5 | 28 | 3047 | 0.005 | 0.116 |
| Metabolic pathways | 60 | 14.9 | 1209 | 3047 | 0.042 | 0.758 |

We attempted to screen the canonical insect’s sex determining genes from our transcriptome, and we then found the single transcript of doublesex, fruitless, and DMRT3, and two transcripts of sex-lethal and transformer-2, however, no transcripts with male- or female-biased expression patterns **(Table S2)**.

## Discussion

To date, RP-HPLC analysis of sinus-gland using several brachyuran species such as the Alaskan tanner crab *Chionoecetes bairdi* and red deep-sea crab *Chaceon quinquedens* have reported two forms of CHH1 and CHH2 as the major form (Chung et al., 2009; Green et al., 2023; Jia et al., 2012). Consistently, the expression level of EiCHHa was 10–100 times higher than CHH1 (EiCHHb and EiCHHc) in this study **(Table 2)**. Moreover, our transcriptome revealed that EiCHHb (Cluster-6453.19920) expression level was significantly higher in females than in males, but not in EiCHHc (Cluster-6453.24714) **(Table 2)**. Although a large dispersion of expression profiles among individuals, the expression level of EiCHHa (Cluster- 6453.16696) seemed to be higher in females as well **(Table 2)**. Moreover, the results of RP-HPLC using sinus gland extraction also demonstrated that the peak of EiCHHa was larger in females **(Figure 2)**. Previous studies reported that the sinus-gland of the same-sized females and males contains similar CHH amount in the European green crab *Carcinus maenas* (Chung and Webster, 1996), however, CHH2 content in the sinus-gland was higher in females than in males of the red deep-sea crab *C. quinquedens* (Green et al., 2023). These data suggested that CHH1 and CHH2 of deep-sea crab species might have female-specific physiological roles. Indeed, the phylogenetic analysis using deduced amino acid sequences of constructed horsehair crab CHHs (EiCHHs) and MIH (EiMIH) with other decapod CHH family showed that female-biased EiCHHb (Cluster-6453.19920) grouped with CHH of snow crab *Chionoecetes opilio* **(Figure 3)**. Likewise, EiMIH grouped with other crab MIHs, implying that EiMIH may have MIH activity. Indeed, our group has also completed the isolation and purification of EiMIH and partial sequence analysis of its amino acid sequence (data not shown).

For some reason, *E. isenbeckii* did not exhibit a decrease in blood glucose levels even after eyestalk ablation followed by fasting. In the control group without eyestalk ablation, blood glucose levels gradually decreased over six days of fasting; however, the extent of this fasting-induced decrease was comparable to that observed in the eyestalk ablation group **(Figure S3)**, indicating that conducting an in vivo assay for EiCHHa using *E. isenbekii* is challenging. Recently, our group demonstrated that an in vivo bioassay using *P. clarkii* is effective for evaluating the physiological activity of CHH purified from the eyestalk of the Japanese spiny lobster *Panulirus japonicus* (Toyota et al., 2024). This method provides a valuable approach for assessing the physiological activity of eyestalk peptides from species that are rare, large, or difficult to use in in vivo assays under laboratory conditions. In this study, the *in vivo* assays of EiCHHa demonstrated its hyperglycemic effect in *P. pelagicus*, a member of the Brachyura, but no such activity was detected in *P. clarkii*, which belongs to the Astacidea. This suggests that functional analysis of the CHH-superfamily is feasible among species within the same infraorder. Furthermore, this finding broadens the potential for *in vivo* analysis of CHH-superfamily peptides isolated from deep-sea, cold-water crustaceans, which are difficult to rear in laboratory settings.

CFSH was identified from the female blue crab *Callinectes sapidus* to develop and maintain secondary female-specific traits, and its expression was dominant in female eyestalk ganglions (Zmora and Chung, 2004). In most decapod species, both CFSH1 and CFSH2 were found, but in the Pacific white shrimp *Litopenaeus vannamei* only three CFSH1 paralogs were found, and no CFSH2 (Veenstra 2016). Here, only one CFSH2 (EiCFSH) was assembled from the eyestalk transcriptome. Although CFSH was initially isolated from eyestalk, in the crayfish *P. clarkii*, its CFSH was strongly expressed in the ovary (Veenstra 2016), indicating that further transcriptome using different tissues of horsehair crabs will be obtained CFSH1. When CFSH was first discovered in the female blue crab *C. sapidus*, it was found as a female sinus gland-specific peptide hormone and showed a female-specific expression pattern in the eyestalk ganglions (Zmora and Chung, 2004). To date, CFSH orthologs have been identified in various crab species such as the swimming crab *Portunus trituberculatus* (Wang et al., 2023), the Chinese mitten crab *Eriocheir sinensis* (Veenstra 2016), and the mud crab *Scylla paramamosain* (Liu et al., 2018), in various shrimp such as the banana shrimp *Fenneropenaeus merguiensis* (Powell et al., 2015), and the kuruma prawn *Marsupenaeus japonicus* (Kotaka et al., 2018). Recent our study has demonstrated that CSFH has the dual roles in *M. japonicus*: growth of male juvenile and regulation of body color by dispersing the pigment granules in the chromatophore (Toyota et al., 2023b). Moreover, the eyestalk transcriptome in the snow crab *C. opilio* revealed that its CFSH was predominantly expressed in the fully matured male with terminal molt (Toyota et al., 2023c). Thus, it is beginning to be pointed out that CFSH may have female- or male-specific roles rather than only female-specific functions. Interestingly, since only EiCFSH was found in the eyestalk of horsehair crab and its expression profile was female-dominant (TPM: 475 in females and 4 in males; **Table 2**), EiCFSH may act on female-specific reproductive organ development and its maintenance as in the blue crab *C. sapidus*.

Of the top ten male-biased transcripts, seven were cuticle-related transcripts, while no cuticle-related DET was found in the female-biased DET list, suggesting that male-specific cuticle proteins are actively transcribed in the eyestalk ganglion of males. A recent study using freshwater prawn *Macrobrachium rosenbergii* revealed that several cuticle-related genes were upregulated during male embryogenesis, suggesting that morphological differences in the cuticle may be constructed in a very early stage of its larval phase (Grinshpan et al., 2022). From another perspective, it is known that there are sex differences in the compound eyes of insects (Zeil, 1983), and that a male-specific gene, *doublesex*, is expressed in the compound eyes of the freshwater crustacean *Daphnia magna* in a male-specific manner (Kato et al., 2011; Toyota et al., 2011). Taken together, eyestalk-compound eye complex contains sexual dimorphisms in various layer of development such as gene expression, compound eye formation, and visual system development. Further study will be required to understand why cuticle-related genes are highly expressed in the eyestalk ganglion of male horsehair crabs.

Recently, many studies have focused on sex determination and sexual differentiation, since fishery-important decapod species including horsehair crabs have generally different values between males and females, for example, growing speed, and maximum body size. Next-generation sequencer-driven transcriptome has enabled to easily obtain expression profiles and comprehensive gene models without a reference genome. Our recent eyestalk transcriptome of kuruma prawn *M. japonicus* revealed no male- or female-specific sinus-gland-derived neuropeptides, however, one of *doublesex* (*MjapDsx1*) shows a higher expression pattern in males than in females (Toyota et al., 2023a). Dublesex is a transcription factor, which is first found in the fruitfly *Drosophila melanogaster* as a master sex regulator (Baker and Wolfner, 1988), and its sex-specific expression is regulated by a series of sex-specific splicing cascades by sex-lethal, transformer, transformer-2, and fruitless (Matson and Zarkower, 2012). In this study, two copies of sex-lethal and transformer-2, and one copy of doublesex and fruitless were predicted, although no these transcripts with male- or female-biased expression patterns **(Table S2)**. This result suggests that male-specific activation of cuticle-related genes is not regulated by *doublesex*, and that sex-specific *doublesex* expression patterns might be divergent among decapod species.

## Conclusion

In this study, transcriptome using male and female eyestalk ganglion has paved the way for understanding the comprehensive gene repertoire of horsehair crab *E. isenbeckii* for the first time. We successfully isolated and purified EiCHHa from sinus gland extraction, determined its full-length sequence through cDNA cloning, and elucidated its hyperglycemic activity through in vivo analysis. Our data showed that two sinus-gland-derived neuropeptides (EiCHHb and EiCFSH) were isolated as female-biased genes, suggesting that both hormones may have female-specific roles such as the development of female characteristics and reproduction. While in the male eyestalk ganglion, many cuticle-related genes were accumulated. In future studies, a substantial functional assay will be necessary to understand the physiological roles of their genes. Although cold-water crustaceans are disadvantaged in physiological experiments, we would like to develop assays using juvenile crabs and their temperate relatives.

## Data availability

Our transcriptome data was deposited in DDBJ Sequence Read Archive (accession number: DRA016047). EiCHHa was also deposited in DDBJ (LC868379).

## Author contributions

Toyota, KT, and TO designed the conception of this study. Toyota, AK, YA, TO conducted the experiment, material preparation, and data collection. Toyota analyzed the RNAseq data. The first draft of the manuscript was written by Toyota and all authors commented on it. All authors read and approved the final manuscript.

## Acknowledgments

This work was supported by the Research Institute of Marine Invertebrates 2019 (KT). Computational resources were provided by the Data Integration and Analysis Facility, National Institute for Basic Biology (Japan).

## Supplemental data

Figure S1

The principal component analysis (PCA) plots of the eyestalk transcriptome analysis (a, b). FF and MM indicate the female and male samples. respectively. Cluster dendrogram of a Euclidean distance, which is computed between samples, and the dendrogram is built upon the Ward criterion (c). The MA-plot of the data for the comparisons done, where differentially expressed features are highlighted in red (d). An MA plot represents the log ratio of differential expression as a function of the mean intensity for each feature. Triangles correspond to features having a too-low/high log2(FC) to be displayed on the plot.

Figure S2

Monitoring of blood glucose levels of male adult *E. isenbeckii* with none-eyestalk ablation (black circle) and with bilateral eyestalk ablation (white circle) for six days.

Figure S3

Maximum likelihood tree of the amino acid sequences of CFSH with 1,000 bootstraps among crustaceans. Accession numbers are in **Table S1**.

Table S1. Sequences of eyestalk-derived peptides used in the phylogenetic analysis

Table S2. Expression profiles of insect’s canonical sex-determining genes

